# A 3’-end capture sequencing method for high-throughput targeted gene expression profiling

**DOI:** 10.1101/2021.12.02.470897

**Authors:** Eric de Bony, Fien Gysens, Nurten Yigit, Jasper Anckaert, Celine Everaert, Eveline Vanden Eynde, Kimberly Verniers, Willem Van Snippenberg, Wim Trypsteen, Pieter Mestdagh

## Abstract

Molecular phenotyping through shallow 3’-end RNA-sequencing workflows is increasingly applied in the context of large-scale chemical or genetic perturbation screens to study disease biology or support drug discovery. While these workflows enable accurate quantification of the most abundant genes, they are less effective for applications that require expression profiling of low abundant transcripts, like long non-coding RNAs (lncRNAs), or selected gene panels. To tackle these issues, we describe a workflow combining 3’-end library preparation with 3’-end hybrid capture probes and shallow RNA-sequencing for cost-effective, targeted quantification of subsets of (low abundant) genes across hundreds to thousands of samples. To assess the performance of the method, we designed a capture probe set for more than 100 mRNA and lncRNA target genes and applied the workflow to a cohort of 360 samples. When compared to standard 3’-end RNA-sequencing, 3’-end capture sequencing resulted in a more than 100-fold enrichment of target gene abundance while conserving relative inter-gene and inter-sample abundances. 3’-end RNA capture sequencing enables accurate targeted gene expression profiling at extremely shallow sequencing depth.

## Introduction

The past decades have witnessed an explosion of high-throughput perturbation screens, in which hundreds to thousands of chemical or genetic perturbations are applied to model cell lines for the purpose of drug or target discovery. Next to simple reporter expression readouts, more complex molecular readouts like transcriptome profiling are increasingly being applied to quantify the phenotypic consequences of such perturbations. This approach, also referred to as molecular phenotyping, requires highly scalable and cost-effective RNA-sequencing workflows to enable transcriptome quantification of hundreds to thousands of samples in a single library preparation. Methods like BRB-Seq^1^ and DRUG-seq^2^ were developed for that purpose and combine sample barcoding during reverse transcription with 3’end-library preparation on pools of 96, 384 or even 1536 samples. Combined with shallow sequencing (1-5 M reads per sample), they enable quantification of 5000-10000 of the most abundant genes for differential gene and pathway expression analyses. While extremely powerful, these methods also present a series of limitations: (i) at shallow sequencing depth, only the most abundant genes are quantified, (ii) abundant genes consist mainly of mRNAs (Figure 1A-B) while some applications may require expression profiling of low abundant genes, like lncRNAs and (iii) some applications may require expression profiling of just a subset of genes representing signatures or pathways of interest. Detection of low abundant genes can be achieved by increasing sequencing depth, but this would negatively impact the per sample cost. On the other hand, if focusing only on gene signatures or specific pathways representing a few hundred genes, most of the sequencing cost is lost to irrelevant genes. RASL-seq applies a targeted probe annealing and ligation strategy for cost effective expression profiling of a few hundred genes^3^. Alternatively, capture sequencing technologies have been described that rely on biotinylated probes to enrich library fragments of interest^4^, typically from total RNA-sequencing libraries.

**Figure 1.**
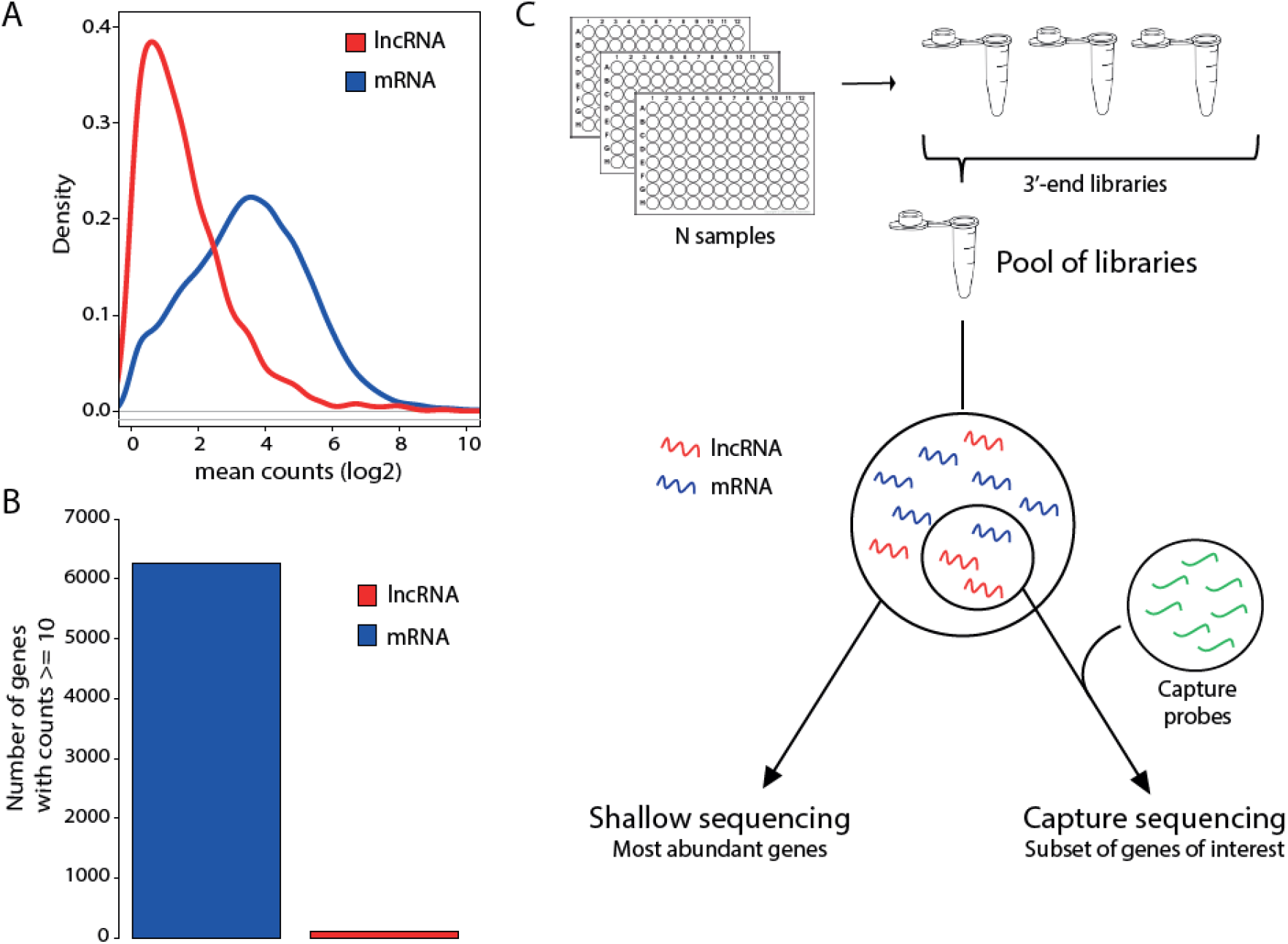
Comparison of the classic 3’-end sequencing workflow and combination with targeted capture sequencing. A. Distribution of mean counts for lncRNAs (red) and mRNAs (blue) by shallow 3’end RNA sequencing across a cohort of 360 samples. B. Number of mRNA (blue) and lncRNA (red) genes with a mean count of at least 10 in the same cohort. C. In the standard 3’end sequencing approach, after introducing a sample barcode at the cDNA synthesis step, libraries are sequenced at low depth resulting in sufficient coverage for the most abundant genes. A subset of (low abundant) genes can be enriched from these pools to avoid reads being consumed by the most abundant genes or simply being allocated to irrelevant genes.

Here, we propose a workflow that combines 3’-end library preparation with 3’-end hybrid capture probes and shallow RNA-sequencing for cost-effective quantification of subsets of (low abundant) genes across hundreds to thousands of samples (Figure 1C). Using a cohort of 360 samples from a CRISPR-interference perturbation screen, we demonstrate efficient and unbiased 3’end capture and sequencing of 137 (mostly low abundant) genes (50 mRNAs and 87 lncRNAs) and compare it to classic shallow 3’end RNA-sequencing.

## Material & methods

### Probe design

Probes were designed using publically available expression data for HEK293T cells and the hg38 genome annotation. For each of the selected genes, probes were designed to target the exon with highest coverage closest to the 3’end (defined based on RNA-sequencing data). For each of these exons, probes of 120 nucleotides were designed, resulting in 3580 probes. Probes were then mapped against repeats and non-targeted genes to remove probes that would capture off-target fragments. Further filtering was done by retaining the 120-mers with a GC content between 25-70%, a GC-based Tm between 60-80 °C and a ΔG larger than −7 (calculated by UNAFold (version 3.8) settings: hybrid-ss-min -E -n DNA -t 54 -T 54). For 25 probes, not all design criteria were met. In total, 482 (Supplementary table 1) probes against 137 unique genes were retained. Probes were synthesized by Twist Biosciences.

### Sample generation and target gene silencing

24 genes were silenced in HEK293T cells using the CRISPR interference (CRISPRi) method^5^. First, nuclease-deficient dCas9-KRAB-MeCP2 (Addgene plasmid no. 110821) was stably introduced in HEK293T cells using the piggy-transposase system (System Biosciences, cat. no. PB210PA-1) according to the manufacturer’s recommendations. dCas9-KRAB-MeCP2-positive HEK293T cells were selected using 10µg/ml of blasticidin. Next, 10 single guide RNAs (sgRNAs) per target were selected from the CRINCL library of sgRNAs. Two negative control sgRNAs (GAACGACTAGTTAGGCGTGTA and GTGCGATGGGGGGGTGGGTAGC) were selected from Horlbeck *et al*.^6^ and Gilbert *et al*.^5^. sgRNA were produced with the Guide-it kit according to manufacturer’s instructions (Takara Bio, cat. nos. 632638, 632639, 632635, 632636 and 632637). Finally, 12,000 cells per well were seeded in 96-well plates (Corning, cat. no. 3596) in 180 µl of RPMI cell culture medium with 10% fetal calf serum. 24 hours after seeding, sgRNAs were transfected with lipofectamine reagent CRISPRMAX (Invitrogen, cat. no. CMAX00003) according to manufacturer’s recommendation. 72 hours after transfection, cells were lysed with SingleShot lysis buffer (Bio-Rad, cat. no. 172-5081).

### 3’end library preparation and 3’-end shallow sequencing

Quantseq RNAseq library preparation (Lexogen) was performed according to the manufacturer’s protocol using 5 µl of cell lysate as input. Libraries were quantified by qPCR, pooled and sequenced on a NextSeq 500 (Illumina). For the reference dataset 428,5M passed filter while 180M reads passed filter for the capture dataset. FASTQ files were processed using an in-house RNA sequencing pipeline. First, FastQC (v0.11.8) was used for data quality control, after which adapter sequences, polyA read-through and low-quality reads were removed via bbduk (BBMap v38.26). Next, reads were mapped with STAR (v2.6.0c, max number of multiple alignments allowed for a read: 20, minimum overhang for unannotated junctions: 8, minimum overhang for annotated junctions: 1, maximum number of mismatches per pair: 999, max number of mismatches per pair relative to read length: 0.1, minimum intron length: 20, maximum intron length: 1000000 and maximum genomic distance between mates: 1000000) against the hg38 reference genome, and gene counts were determined via HTSeq (v.0.11.0).

### Library capture and sequencing

First, 115.3ng of the Quantseq library pool was dried using vacuum centrifuge with no heat. Custom probes were resuspended in 10mM TE buffer (pH 8.0) then mixed with water and TWIST hybridization mix according to the manufacturer’s recommendations. Probe solution was then heated at 95°C for 2 minutes, cooled on ice for 5 minutes then equilibrated to room temperature for 5 minutes. Dried pool was combined with 5µl blocker solution and 8 µl of universal blockers (TWIST Bioscience). Pool solution was then heated at 95°C for 5 minutes then equilibrated at room temperature for no more than 5 minutes. Capture probe solution was then added to the pool solution, solutions were homogenized by pipetting up and down gently 15 times with caution not to generate bubbles. Finally, probe-pool mix were incubated at 70°C for 16 hours in a thermal cycler with lid at 85°C. The next day, 100µl of streptavidin beads (ThermoScientific) were washed 3 times in 200µl of binding buffer. Hybridization reaction was then added to 200µl of washed beads and incubated for 30 minutes at room temperature with agitation. Beads clean-up was then performed according to the manufacturer’s recommendations. After clean-up, 22.5µl of bead slurry was combined with 2.5µl of oligo-dT primers and 25µl of KAPA HiFi HotStart Ready Mix (Sigma Aldrich) and subjected to PCR amplification on a thermal cycler. PCR program: 98°C for 45 seconds; followed by 14 cycles of 98°C for 15 seconds, 60°C for 30 seconds and 72°C for 30 seconds; 72°C for 1 minute then cooled to 4°C. Cool amplified products were then subjected to AMPure beads clean-up according to recommendations. Library was quantified by qPCR and sequenced on a NextSeq 500 (Illumina). FASTQ files were processed using the same pipeline as described above.

### Data analysis

Gene counts were further processed with R (v.4.0.3). Samples with less than 100000 counts were not considered for further data analysis. Gene counts were normalized to Reads Per Million (RPM). Gene coverage plots were created using Sushi package (Sushi_1.28.0) in R studio. When comparing expression of sgRNA targets between datasets, only targets with a mean count of 5 or higher in the reference dataset were considered. Log2 of gene expression fold-changes were calculated using RPMs. Simulated sequencing depths required to reach minimal coverage for a random subset of 100 genes were calculated by summing reads required to reach said coverage for the least abundant gene and 99 others (for capture) and all other detected genes (without capture).

## Results

### 3’-end capture efficiently enriches low abundant targeted isoforms

To evaluate our shallow 3’-end capture sequencing workflow, we used a collection of 360 samples from HEK293T cells stably expressing the dCas9-KRAB-MeCP2 protein. These cells were treated with individual sgRNAs to silence various mRNAs and lncRNAs with CRISPR-interference (see Methods). A pool of 500 3’end capture probes were designed to target 137 genes (50 mRNAs and 87 lncRNAs, including those targeted by the sgRNAs). Target genes were distributed across abundance spectrum with 4.4% presenting less than 1 RPM, 50.3% with less than 10 RPMs, 93.4% with less than 100 RPMs, 99.3% with less than 1000 RPMs and 100% with less than 10000 RPMs. We will further refer to these genes as the captured genes. Each 3’-end was covered with at least one and up to four 120 bp capture probes. To establish a reference gene expression profile for these samples, we applied 3’end library preparation and shallow sequencing to all samples (we will refer to this as the reference dataset). An aliquot of the pooled 3’-end libraries was subsequently subjected to hybrid capture in a single-tube reaction and was sequenced at similar coverage as the reference. In the reference dataset, 35/137 captured genes were detected with a mean of at least 10 counts. The majority was either not detected (had no counts, n = 16) or had between one and 10 counts (n = 86). With capture, 123/137 captured genes had at least 10 counts, 10 had between 1 and 10 counts and only four had less than one count. Sequencing at a 10-fold lower depth would still have resulted in 100/137 captured genes with 10 or more counts, suggesting that 3’end capture sequencing enables a considerable reduction of sequencing depth with limited impact on target representation. The capture approach resulted in a mean enrichment of 255-fold (CI 95%: 204-306) when comparing reads per million (RPM) of the captured genes between reference and capture datasets. The enrichment was similar for both lncRNAs and mRNAs (Figure 2A). The fraction of reads assigned to the capture genes increased from 1.2% in the reference to 78.4% in the capture dataset (Figure 2B). This was accompanied by a significant reduction in library complexity and a shift of captured genes from being among the least abundant genes in the reference dataset to the most abundant genes in the capture dataset (Figure 2C). In total, capture was not successful for 16 out of 137 genes. For these genes, capture either failed to increase the mean gene count to 10 or more (14 genes) or capture failed to induce a two-fold enrichment for genes with a mean count of 10 or higher (2 genes). In all these cases, inspection of the 3’-end coverage revealed that the capture probes did not overlap the 3’end library fragment.

**Figure 2.**
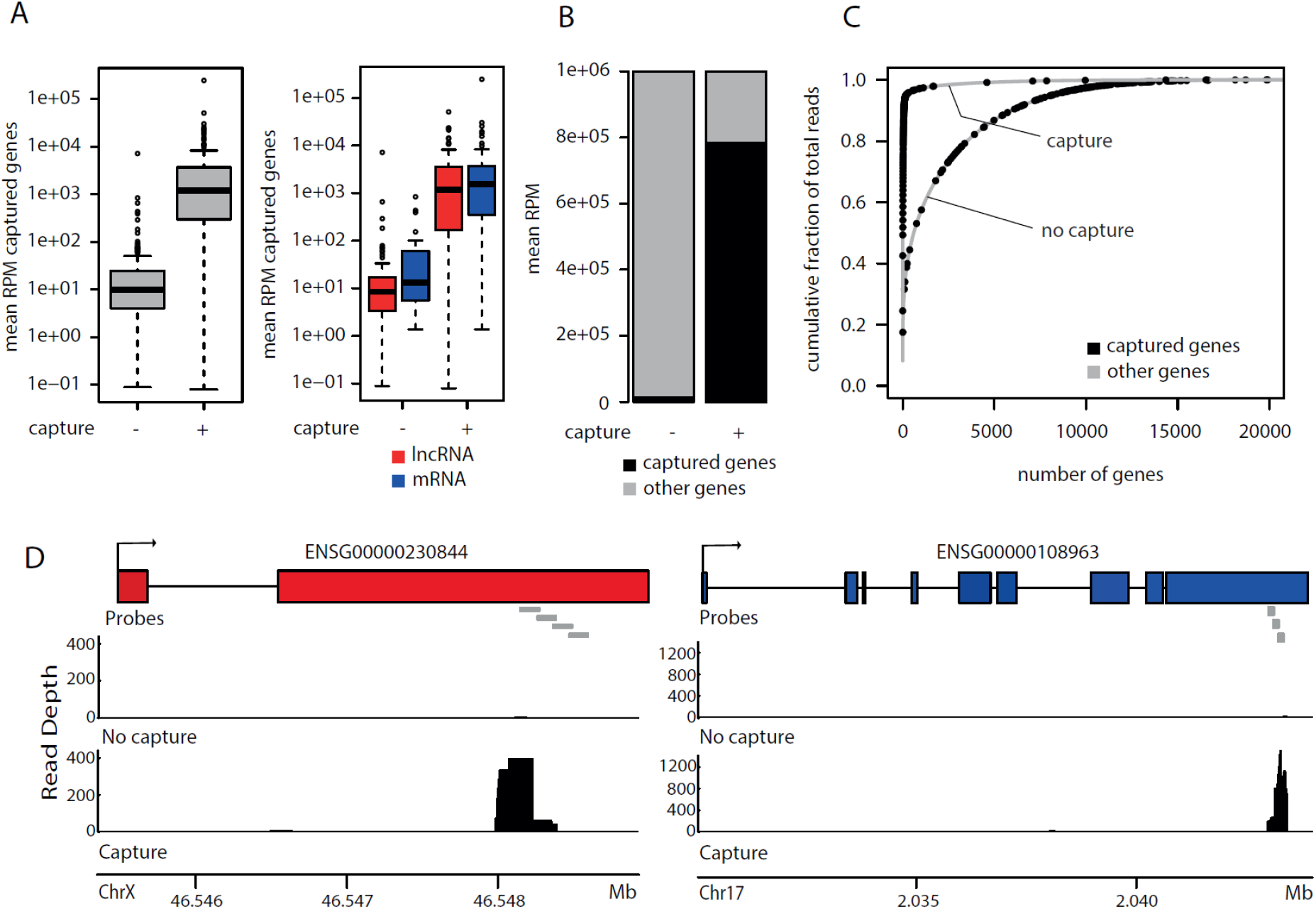
Capture sequencing reveals similar enrichment for mRNAs and lncRNAs. A. Mean RPM enrichment after capture for 87 lncRNAs and 50 mRNAs. B. Read allocation for the 137 targeted genes with or without capture. C. Cumulative fraction of total reads with or without capture. D. Coverage plot for a lncRNA (red) and mRNA (blue).

As expected, capture results in a coverage increase at the 3’end of the capture genes, coinciding with the position of the capture probes (Figure 2D). Except for the genes for which capture was unsuccessful, capture efficiency was uniform across samples and across captured genes (Figure S1), independently from initial abundance (Figure S2). Taken together, these results demonstrate that 3’end library preparation combined with hybrid capture can be applied to enrich low abundant genes that are not detectable or poorly covered in a standard shallow 3’-end RNA-sequencing workflow.

### Capture maintains inter-gene and inter-sample gene abundance differences

To further validate the quantitative performance of the 3’end capture sequencing workflow, we first compared the abundance of captured genes between the reference and capture datasets. We observed a significant correlation of RPM values of captured genes between the reference and capture dataset (r = 0.492, p = 1.01e-9, spearman), suggesting that inter-gene abundance differences are conserved for the majority of captured genes (Figure 3A). As our dataset contained samples treated with negative control sgRNAs and sgRNAs against specific mRNA and lncRNA targets, we evaluated inter-sample abundances by comparing fold changes for 9 of these targets between the reference and capture dataset. Fold changes were positively correlated (r = 0.9, p = 0.002, spearman) (Figure 3B) and sgRNA target knockdown was conserved between the reference and capture datasets (Figure 3C). These results further confirm the quantitative performance of the 3’end capture sequencing workflow.

**Figure 3.**
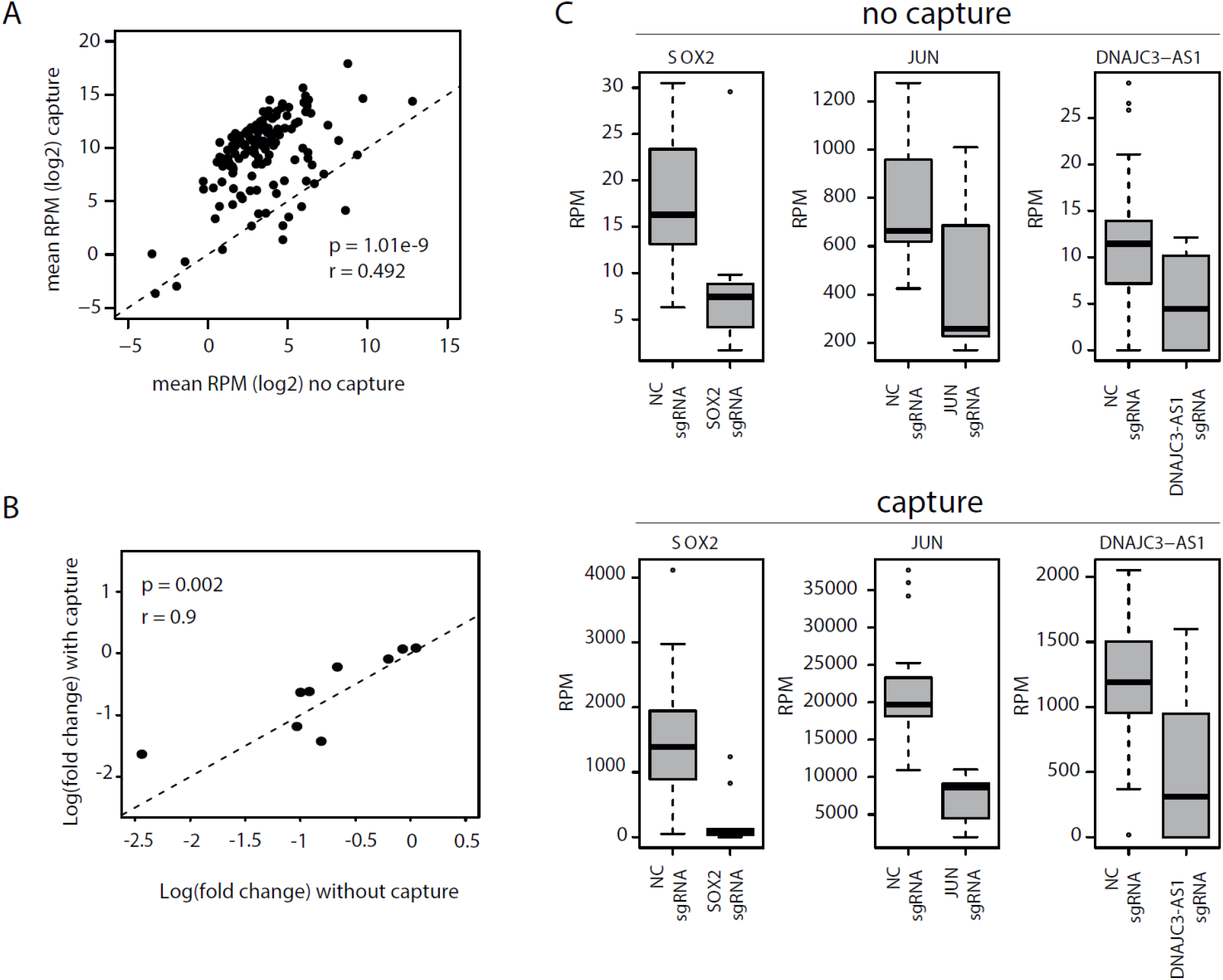
Capture sequencing conserves inter-samples gene relationships. A. Correlation between RPM for captured genes with or without capture. B. Correlation between observed expression fold-changes between sgRNA and NC treated conditions with and without capture for genes presenting a mean count of at least 5 without capture. C. Top, expression in sgRNA treated cells and negative control (NC) treated cells for 3 captured genes undergoing CRISPRi-based silencing in 3’end shallow sequencing. Bottom, same but after capture sequencing.

### Targeted 3’-end capture enables a 100-fold sequencing depth reduction

To further demonstrate the value of our approach, we created a gene panel by randomly selecting 100 genes from the 3’-end shallow sequencing reference dataset and calculated sequencing depths required to achieve a minimal coverage for each gene in that set when applying capture sequencing. Simulations were run based on the observation that, after capture, 80% of all reads are allocated to captured genes (Figure 2B). We observed that the required sequencing depth to reach a given minimal coverage is 100-fold lower when applying capture compared to standard 3’-end sequencing (Figure S3).

## Discussion

For large sample cohorts (hundreds or thousands of samples), 3’-end library preparation followed by shallow sequencing represents a cost-effective solution to monitor the expression of 5000 to 10000 of the most abundant genes. This approach provides a good representation of transcriptomic response to a given perturbation (chemical or genetic). However, large cohort screenings may focus on a subset of genes belonging to a same pathway or signal transduction network, or to a class of low abundant genes like lncRNAs. For such applications, we demonstrate that 3’-end shallow RNA-sequencing approaches can be augmented with hybrid capture. Capture maximizes the allocation of sequencing reads to genes of interest while substantially reducing required sequencing depth. Capture efficiency was similar for coding and non-coding genes and ensured that biological insights such gene expression differences between samples was conserved. We calculated that a capture step, which is performed on the pool of 3’end libraries in a single tube reaction, results in a 100-fold reduction in required sequencing depth. For a set of 100 randomly selected genes, where one aims to generate at least 10 counts per gene (i.e. 10 counts on the least abundant gene), 3’end capture sequencing would require a total of 10000 reads per sample while regular 3’end sequencing would require 1,000,000 reads per sample. On a cohort of 4000 samples, the difference would be 1 NextSeq 500 High Output run with capture versus 10 such runs without capture, and thus a substantial difference in the per sample cost. For low abundant genes like lncRNAs, which are emerging as novel players in all aspects of cellular physiology and disease, shallow 3’-end RNA-sequencing lacks sensitivity. Hybrid capture can enrich lncRNA 3’end fragments, enabling accurate lncRNA expression quantification, even at (extremely) shallow sequencing depth. Given proper probe design, high and consistent capture efficiency can be obtained. While capture failed for 16/137 targeted genes, closer inspection of the capture probe location for these genes revealed that these did not overlap the regions with 3’end coverage, but were positioned just up-or downstream of the 3’end coverage peak. We used publicly available total RNA-sequencing data of the studied cell line to define 3’ends of the target genes for probe design. As total RNA-seq coverage is known to be less accurate towards 5’ and 3’ transcript boundaries, probe design is best informed using actual 3’end RNA-sequencing data from the cell line under investigation. Of note, stringency of probe design criteria can potentially be reduced, as 22 low quality, yet correctly positioned, probes lead to successful enrichment (supplemental table 1). Taken together our results demonstrate the value of combining 3’-end library preparation with hybrid capture to enable highly scalable and cost-effective expression profiling of (low abundant) genes of interest.

## Acknowledgements

This work was supported by the European Union’s Horizon 2020 research and innovation program under grant agreement 826121, the and the Concerted Research Action of Ghent University (BOF/GOA 01G00819) and the Fund for Scientific Research Flanders (FWO).

## References

1. Alpern, D. et al. BRB-seq: ultra-affordable high-throughput transcriptomics enabled by bulk RNA barcoding and sequencing. Genome Biol. 20, 71 (2019).

2. Ye, C. et al. DRUG-seq for miniaturized high-throughput transcriptome profiling in drug discovery. Nat. Commun. 9, 4307 (2018).

3. Li, H., Qiu, J. & Fu, X.-D. RASL-seq for massively parallel and quantitative analysis of gene expression. Curr. Protoc. Mol. Biol. Chapter 4, Unit 4.13.1-9 (2012).

4. Mercer, T. R. et al. Targeted sequencing for gene discovery and quantification using RNA CaptureSeq. Nat. Protoc. 9, 989–1009 (2014).

5. Gilbert, L. A. et al. CRISPR-Mediated Modular RNA-Guided Regulation of Transcription in Eukaryotes. Cell 154, 442–451 (2013).

6. Horlbeck, M. A. et al. Compact and highly active next-generation libraries for CRISPR-mediated gene repression and activation. eLife 5, (2016).

