## Supplementary material for "A 3’-end capture sequencing method for high-throughput targeted gene expression profiling": Supplmental figure 1-3

**Supplemental table 1.** 500 TWIST probes against 137 unigque genes to perform capture sequencing.

**Supplemental figure 1.** The ratio between RPM's for the 100 most abundant capture genes.

**Supplemental figure 2.** The initial abundance (mean RPM before capture) compared to the capture efficiency (mean RPM ratio)

**Supplemental figure 3.** Simulation of the needed sequencing depth for a random set of a 100 genes of interest.

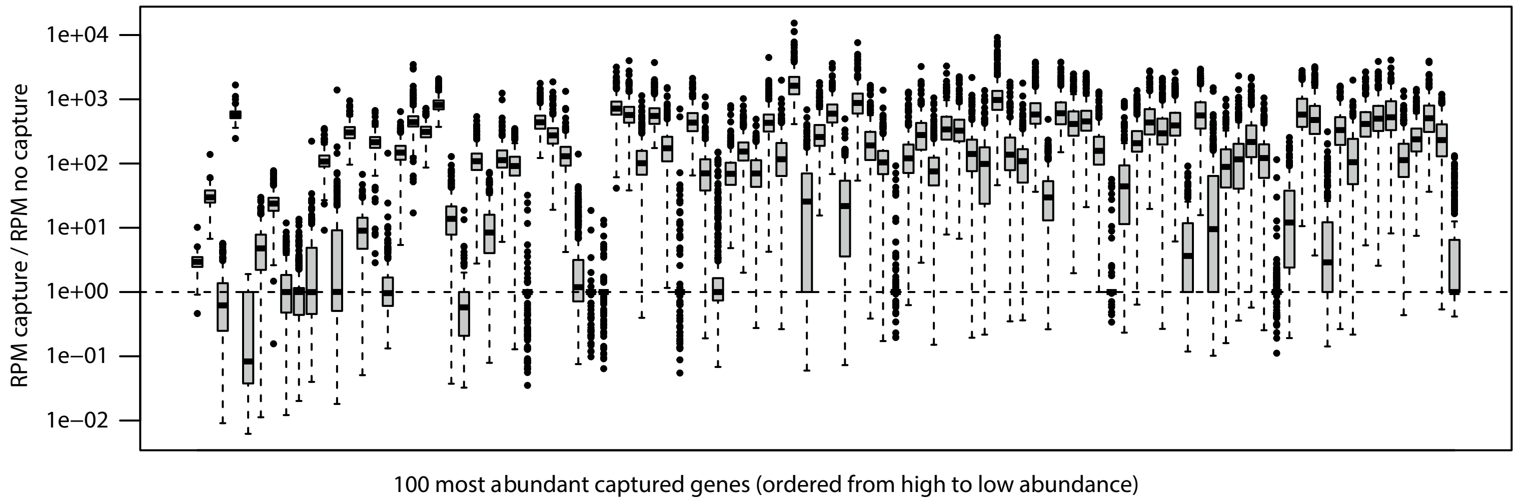
1e+04
1e+03
1e+02
1e+01
1e+00
1e−01
1e−02
RPM capture / RPM no capture
100 most abundant captured genes (ordered from high to low abundance)

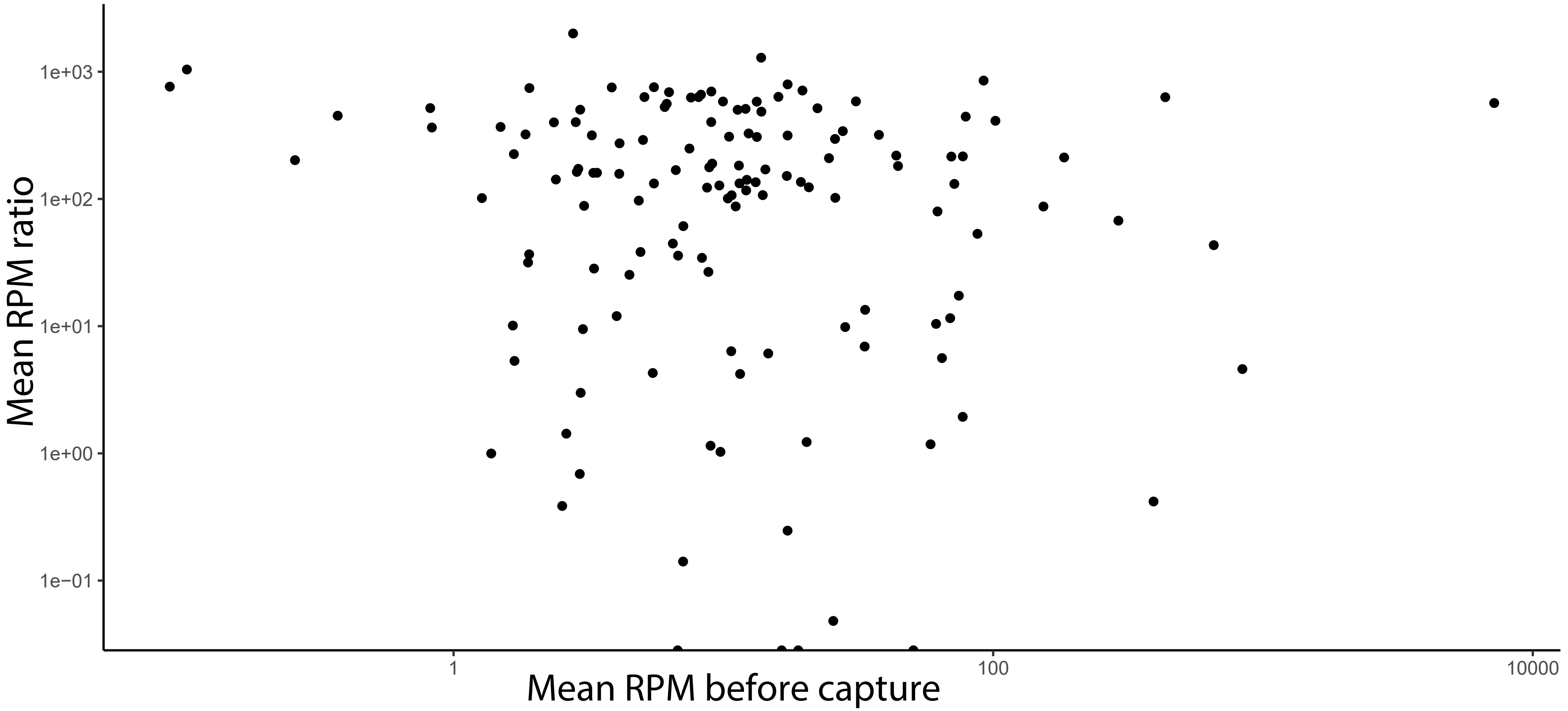
1e+03
1e+02
1e+01
1e+00
1e−01
Mean RPM ratio
1
100
10000
Mean RPM before capture

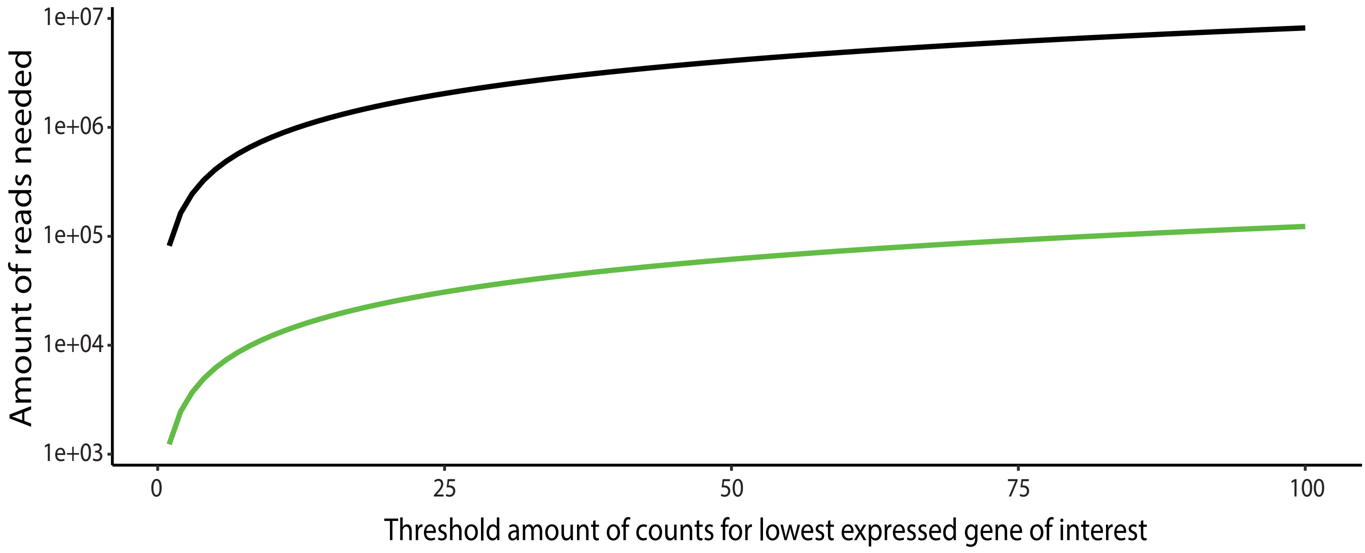

Amount of reads needed
1e+07
1e+06
1e+05
1e+04
1e+03
0
25
50
75
100
Threshold amount of counts for lowest expressed gene of interest
